# Kilohertz two-photon fluorescence microscopy imaging of neural activity *in vivo*

**DOI:** 10.1101/543058

**Authors:** Jianglai Wu, Yajie Liang, Shuo Chen, Ching-Lung Hsu, Mariya Chavarha, Stephen W Evans, Donqging Shi, Michael Z Lin, Kevin K Tsia, Na Ji

## Abstract

Understanding information processing in the brain requires us to monitor neural activity *in vivo* at high spatiotemporal resolution. Using an ultrafast two-photon fluorescence microscope (2PFM) empowered by all-optical laser scanning, we imaged neural activity *in vivo* at up to 3,000 frames per second and submicron spatial resolution. This ultrafast imaging method enabled monitoring of both supra- and sub-threshold electrical activity down to 345 μm below the brain surface in head fixed awake mice.

---

The ability to monitor neural signaling at synaptic and cellular resolution *in vivo* holds the key to dissecting the complex mechanisms of neural activity in intact brains of behaving animals. The past decade has witnessed a proliferation of genetically encoded fluorescence indicators that monitor diverse neural signaling events *in vivo*, including those sensing calcium transients, neurotransmitter and neuromodulator release, and membrane voltage [1]. Most popular are the calcium indicators (e.g., GCaMP6 [2]) and glutamate sensors (e.g., iGluSnFR [3]), with their success partly attributable to their slow temporal dynamics (e.g., rise and decay times of 100 – 1000’s milliseconds for GCaMP6 and tens of milliseconds for iGluSnFR), which can be adequately sampled with conventional 2PFM systems. Indeed, using point scanning and near-infrared wavelength for fluorescence excitation, 2PFM can routinely image calcium activity hundreds of microns deep in opaque brains with submicron spatial resolution [4–6].

Imaging faster events, however, is more challenging. Indicators reporting membrane voltage, arguably the most direct and important measure of neural activity, have rise and decay times measured in milliseconds. Too fast for the frame rate of conventional 2PFM to match, their *in vivo* imaging demonstrations were mostly carried out by widefield fluorescence microscopy with comparatively poor spatial resolution and limited to superficial depths of the brain [7–9]. In other words, the capability of state-of-the-art indicators has outstripped our ability to image them at sufficiently high speed, especially at high spatial resolution and through highly scattering brain tissue.

To observe millisecond dynamics such as membrane potential variations, we need to increase 2PFM frame rates to the kHz range. The sampling rate of conventional raster-scanning 2PFM is limited by the speed of laser scanners, such as galvanometric mirrors, to tens of frames per second (fps) [5, 6]. Random-access 2PFM using acoustic-optical deflectors allows kHz frame rates and has been used to monitor the membrane potential of neurons expressing genetically encoded voltage indicators (GEVIs) in brain slices and *in vivo* [10, 11], but can only track a pre-selected set of locations (currently up to ∼20). A line-projection tomographical 2PFM system was used to detect glutamate release in the brain at kHz frame rates [12], but is limited in the complexity of fluorophore distributions that can be reconstructed for a given number of excitation angles. Notably, in these 2PFM implementations, the pixel dwell times (0.1 – 40 microseconds) are much longer than the theoretical minimum dwell time required for assigning signals to scanned locations. This minimum dwell time is on the order of several nanoseconds, limited only by the fact that dwell times shorter than fluorophore excited state lifetimes would allow emission photons to return from multiple excited locations at the same time [13]. Here, by leveraging an all-optical passive laser scanner based on a concept termed free-space angular-chirp-enhanced delay (FACED) [14], we demonstrated raster-scanning 2PFM at 1,000 fps and 3,000 fps, with the pixel dwell time flexibly configured to reach the fluorescence lifetime limit. We applied it to ultrafast monitoring of calcium activity, glutamate release, and membrane potential with a variety of genetically encoded activity indicators, and demonstrated ultrafast 2PFM imaging of spontaneous and sensory-evoked supra- and sub-threshold electrical activity in awake mouse brains *in vivo*.

The principle of FACED was detailed previously (**Fig. 1a**) [14]. Briefly, a pulsed laser beam was focused in 1D by a cylindrical lens and obtained a converging angle *Δθ*. It was then launched into a pair of almost parallel high-reflectivity mirrors with separation *S* and misalignment angle *α*. After multiple reflections by the mirrors, the laser beam/pulse was split into multiple beamlets/subpulses (*N* = *Δθ/α*) of distinct propagation directions and eventually retroreflected with an inter-pulse temporal delay of 2*S/c*, with *c* being the speed of light. After being relayed to enter a microscope objective, this pulse train formed an array of spatially separated and temporally delayed foci (**Fig. 1b**). In a fluorescent sample, they excited two-photon fluorescence in succession, which could be detected by a photomultiplier tube, sampled at high speed, and assigned to individual foci and image pixels. Effectively, this passive FACED module allows line scanning at the repetition rate of the pulsed laser, typically MHz.

**Figure 1:**
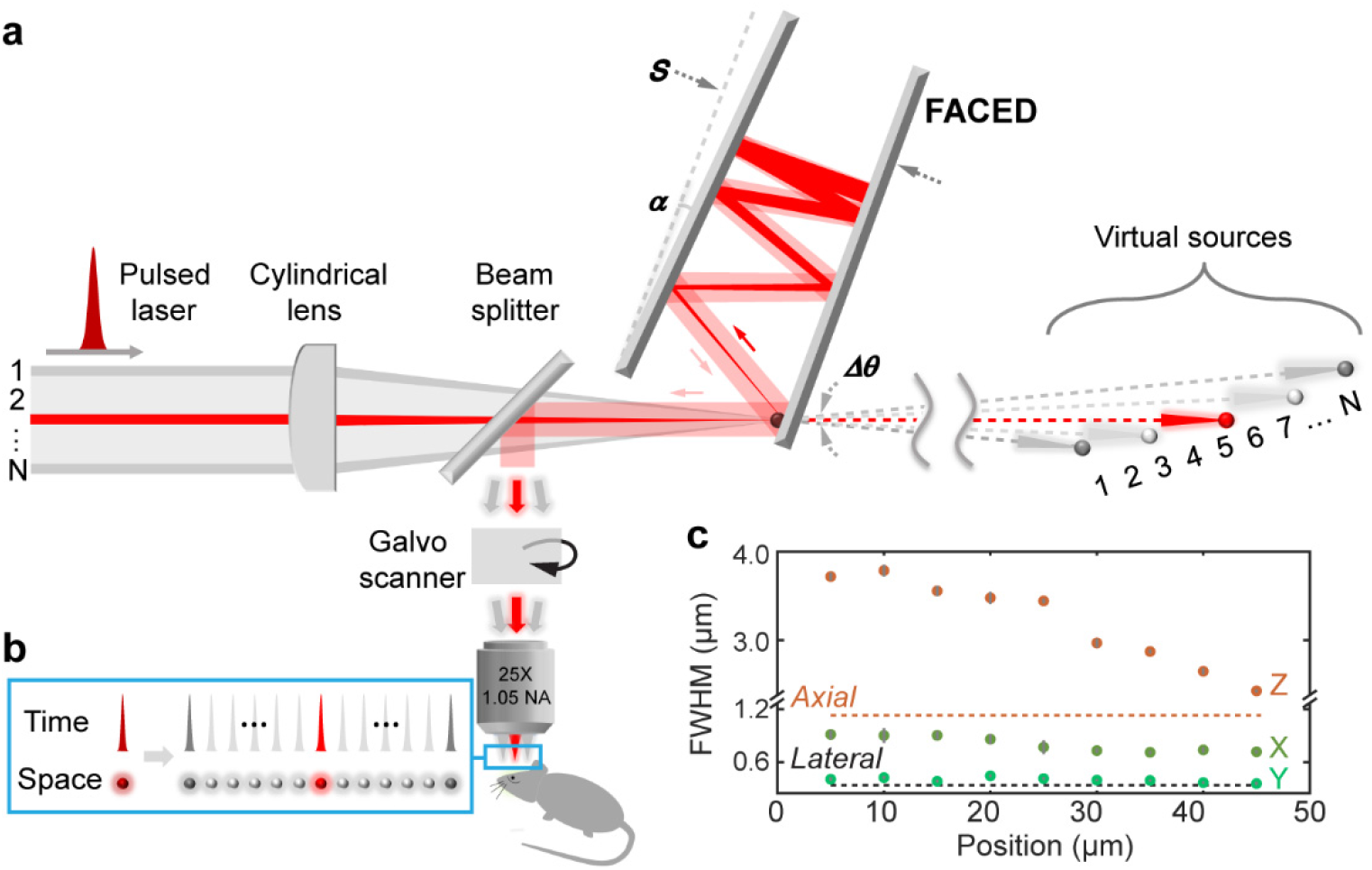
Principles and resolution of a 2PFM with a FACED module. (a) Schematic of a FACED microscope. A 1-MHz collimated femtosecond laser was focused into a nearly parallel mirror pair with a converging angle *Δθ* by a cylindrical lens. After multiple reflections, the misalignment angle *α* caused the beamlets to retroreflect (e.g., the red rays). Beamlets at different incidence angles (e.g., red versus gray rays) emerged with distinct propagation directions and temporal delays. Equivalently, the sequence of multiple beamlets (*N* = *Δθ/α*) at the output of the FACED module can be treated as light emanating from an array of virtual sources. These beamlets were then coupled into a 2PFM and formed (b) an array of spatially separated and temporally delayed foci at the focal plane of a microscope objective. (c) The focal spot sizes along the X/FACED, Y, and Z axes, measured from 200-nm-diameter fluorescent beads. Error bars show s.d. from 10 beads; dashed lines indicate the expected axial and lateral resolutions at 1.05 NA.

In this work, we designed a FACED module and incorporated it into a standard 2PFM upgraded with a high-speed data acquisition system (625 MS/s) (**Supplementary Methods**, **Supplementary Fig. 1**). Using a 920-nm laser with a 1-MHz repetition rate, our FACED module produced a line of 80 pulsed foci spanning 50 μm each microsecond. The inter-pulse interval was set at 2 ns to reduce pixel crosstalk due to fluorescence lifetime and detector response time. We measured the full width at half maxima (FWHMs) of these foci by imaging 200-nm-diameter fluorescent beads. From the first to last, the foci were images of virtual sources formed after distinct numbers of mirror reflections and located at increasing distances away (**Fig. 1a**), with the more distant virtual sources leading to larger beam sizes at the back focal plane of the objective. As a result, the more temporally delayed pulses had smaller foci along both the lateral and axial directions (**Fig. 1c**). The beams filled the back focal plane more along the Y than X/FACED axis, resulting in ∼0.82 μm (X) and ∼0.35 μm (Y) lateral resolution, sufficient to resolve subcellular structures. With the FACED module providing an 80-pixel line each microsecond, we scanned an entire frame of 80 × 900 pixels at 1,000 Hz by scanning the Y galvanometer at 500 Hz and collecting data bidirectionally. An additional X galvanometer allowed us to tile the images to cover a larger field of view, if desired.

We first used the FACED 2PFM to image calcium dynamics using genetically encoded calcium indicators GCaMP6s and 6f [2]. At 1 kHz, morphological features were clearly resolved in individual images (see **Supplementary Fig. 2** for representative raw images taken at 1 kHz). We reliably detected calcium transients in GCaMP6s-expressing cultured neurons (**Supplementary Fig. 3** and **Video 1**) and GCaMP6f-expressing acute mouse brain slices (**Supplementary Fig. 4** and **Video 2**) that were evoked by extracellular electric stimulation. Due to its high spatiotemporal resolution, we could clearly resolve neurites in cultured neurons, and in one case, we recorded spontaneous calcium releases in neurites which then propagated at 25 μm/s across the dendrites (**Supplementary Fig. 5** and **video 3**).

Clearly, the true power of FACED lies in imaging faster physiological events. Next, we imaged neurons labeled with the A184V (medium-affinity) and S72A (low-affinity) variants of the genetically encoded glutamate sensor SF-iGluSnFR, which are expressed on cell membranes and report spikes with faster activation and inactivation kinetics than the calcium indicators [15]. FACED 2PFM reliably reported glutamate release events evoked by field stimulation in cultured neurons (**Fig. 2a** and **Supplementary videos 4-7**), as well as spontaneous glutamate release in L2/3 neurons in the primary visual cortex (V1) of head-fixed awake mice (**Fig. 2b** and **Supplementary videos 8-9**). Both in culture and *in vivo*, we observed faster dynamics from the lower-affinity variant S72A than A184V, consistent with their previous characterization by conventional 2PFM [15]. Imaging the brain *in vivo* at 1,000 fps, we also observed rapid movements of fluorescent particles, which transited the field of view at ∼1 mm/s (**Supplementary Fig. 6** and **Supplementary video 10**). We speculated that they were macrophages containing fluorescent remnants of dead cells, moving rapidly with blood flow in the vasculature. The ability of FACED 2PFM to capture such rapid events indicates that this method can also be used to study rapid biological events associated with blood flow.

**Figure 2:**
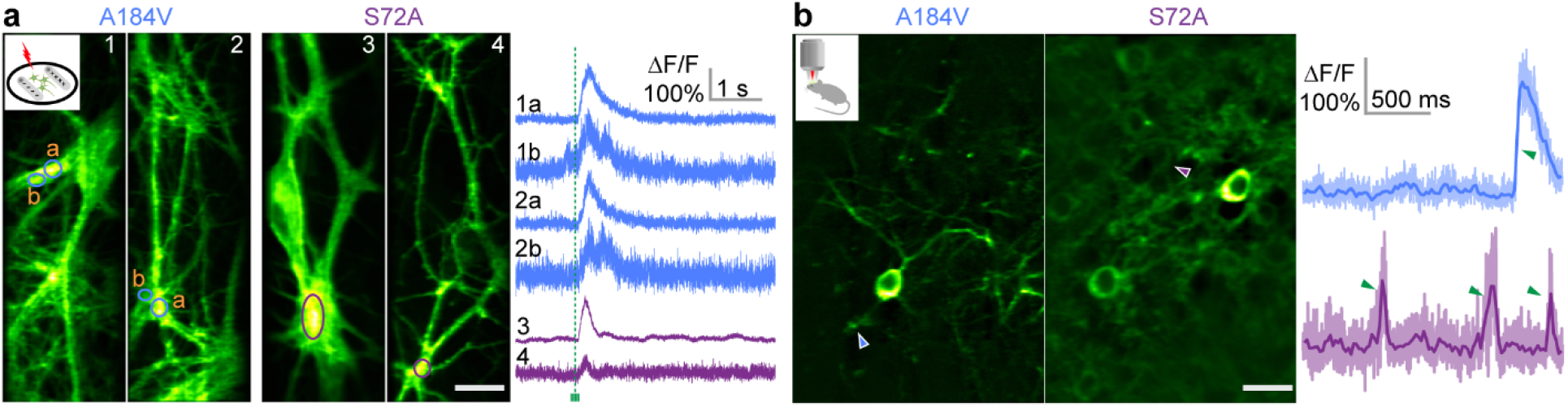
1 kHz imaging of genetically encoded glutamate indicator iGluSnFR variants in cultured neurons and in V1 of awake mice *in vivo*. (a) (Left) mean intensity projections of cultured neurons expressing either the A184V or the S72A variants of SF-iGluSnFR; (Right) transients associated with glutamate release triggered by extracellular electric stimulation (green dashed line) for structures labeled on the left. (b) (Left) Representative images of layer 2/3 neurons (depth: 150 – 250 μm) expressing either A184V or S72A in V1 of awake mice; (Right) Transients associated with spontaneous glutamate releases at sites indicated by arrowheads on the left. Here, darker lines were 20-point boxcar averages of the raw traces to guide the eye. Green arrowheads on the traces highlight the fast rising edge of the glutamate transients. Scale bars: 20 μm. “Green hot” lookup table in ImageJ was applied to all images. Post-objective power: 25 – 30 mW for cultured neurons, 40 mW *in vivo*.

Finally and most importantly, we imaged neurons expressing the GEVI ASAP3 [10] in V1 of head-fixed awake mice. Among GEVIs, ASAP-family indicators are currently the only ones to have reported single spikes *in vivo* with 2PFM, albeit in a limited number of neurons in the context of random-access scanning [10, 11, 16]. ASAP3 reports membrane depolarizations and action potentials as downward deflections in fluorescence. Using the soma-targeted ASAP3-Kv in both sparsely labeled (**Fig. 3a**) and densely labeled preparations (**Fig. 3b**), we observed that fluorescence was largely concentrated to somata. Imaging these neurons at 1,000 fps continuously for six seconds, we observed substantial photobleaching (**Supplementary Fig. 7**). However, if 1-kHz imaging was carried out in 1-s bouts interleaved with 2.5-s dark periods, ASAP3 fluorescence from individual neurons recovered almost completely between bouts (**Supplementary Fig. 8**), allowing us to interrogate voltage responses from the same neurons repeatedly. Here, we employed both 6-s continuous recordings and intermittent 1-s recordings for *in vivo* experiments.

**Figure 3:**
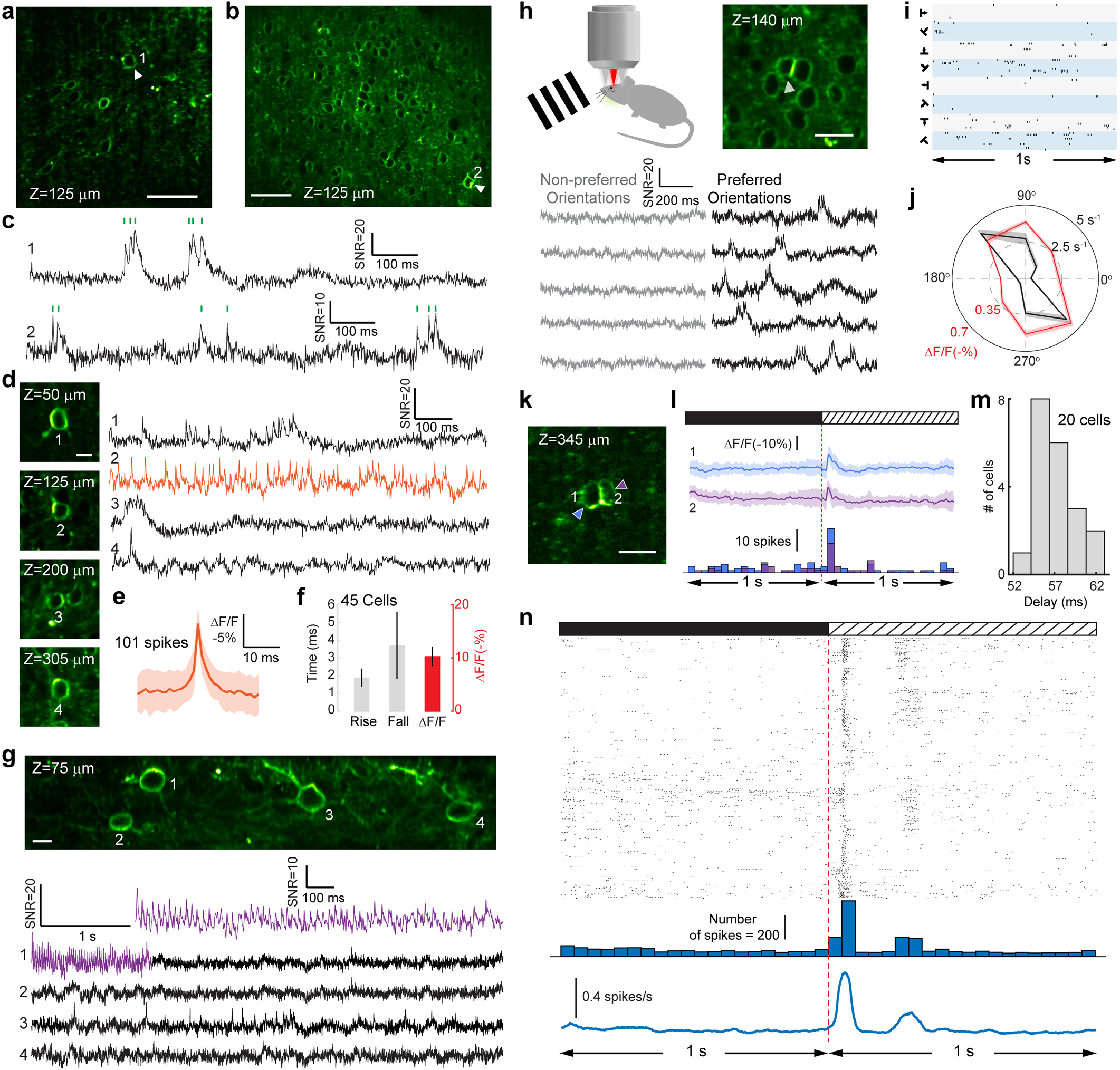
1 kHz imaging of supra- and sub-threshold voltage responses with genetically encoded voltage sensor ASAP3 in V1 of awake mice. (a,b) Representative images of neurons in V1 (a) sparsely or (b) densely labeled with soma-targeted ASAP3-Kv. (c) Spontaneous voltage traces (SNR, ΔF/√F) from neurons 1 and 2 in (a) and (b), respectively; Green ticks: spikes. (d) Neurons at different depths of a cortical column and their spontaneous voltage traces. (e) Average of 101 spikes from neuron 2 (orange trace) in (d); (f) Rise time, fall time, and ΔF/F of spikes (mean ± s.d., n=45 cells from 3 mice). (g) 1kHz imaging over a 50 × 250 μm^2^ field of view with four neurons exhibiting distinct spontaneous activity patterns. (h) Voltage traces from a V1 neurons showing orientation selectivity, with more sub- and supra-threshold activity for preferred orientations (black traces) than for non-preferred orientations (gray traces). (i) Raster plot of spikes for all trials (10 trials for each of 8 grating stimuli) and (j) polar plot showing the orientation tuning of mean subthreshold ΔF/F response (red) and spiking rate (black) of the neuron in (h). (k) Image of two neurons at 345 μm depth and (l) their subthreshold ΔF/F (upper traces, average of 52 trials) and spiking (lower histograms, 50-ms bins) responses relative to the onset of grating stimuli (red dashed line). (m) Histogram of the time to reach peak subthreshold voltage responses post stimulus onset from 20 cells in 3 mice. (n) Spiking response relative to stimulus onset. From top to bottom: raster plot, histogram (50-ms bins), and averaged firing rate (50-ms sliding rectangular windows) of 2747 detected spikes from 617 trials of 20 neurons in 3 mice. Shaded areas: s.d.; Scale bars: (a,b) 50 μm; (d,g) 10 μm; (h, k) 20 μm.

We extracted fluorescence from pixels representing cell membranes to obtain time-dependent traces for each neuron. Despite photobleaching, downward signals corresponding to putative individual action potentials (“optical spikes”) could be easily detected from raw traces in single trials (**Supplementary Fig. 9**). After correcting for photobleaching, we calculated the relative fluorescence change both in terms of signal-to-noise ratio (SNR, ΔF/√F) and ΔF/F (**Methods**, **Supplementary Fig. 9, Fig. 3**).

FACED 2PFM enabled us to detect both suprathreshold and subthreshold membrane potential variations from neurons in both sparsely and densely labeled brains (**Fig. 3c**). The ability of 2PFM to provide high-resolution images in optically opaque brains enabled us to measure spontaneous voltage activity from neurons located at four different depths down to 305 μm below the dura in the same awake mouse brain (**Fig. 3d**). In one of these neurons (orange trace, ROI2 of **Fig. 3d**), we detected a total of 101 isolated spikes in 48 s of recording and obtained an averaged spiking profile (**Fig. 3e**). From the averaged spike waveforms of this and other neurons, we determined the rise time, the decay time, and the ΔF/F for single action potentials to be 1.93 ± 0.51 ms, 3.73 ± 1.88 ms, and 0.10 ± 0.02 (mean ± s.d., n = 45 cells from three mice), respectively (**Fig. 3f**). These voltage response characteristics are consistent with those measured from ASAP3 by random-access two-photon microscopy [10].

FACED 2PFM can image a large field of view (FOV) along the Y-axis without compromising imaging speed. In one example (**Fig. 3g**), we measured both spontaneous spiking and subthreshold activity from four neurons within a 50 × 250 μm^2^ FOV, and observed spike bursts lasting over 1 s in one neuron (purple trace, ROI1 of **Fig. 3g**). Because two-photon excitation is restricted to a femtoliter volume at the focal point and fluorescence photons generated at different locations are detected at different times, there was no cross-pixel contamination of voltage signals even in immediately neighboring neurons (for an example of densely labeled brains, see **Supplementary Fig. 10**). In many recording sessions, we observed rhythmic subthreshold oscillations in the voltage traces of individual neurons (**Supplementary Fig. 11**), with the dominant oscillation frequencies falling within 4-12 Hz, suggesting that they originated from cortical theta oscillations [17, 18].

FACED 2PFM also detected sensory evoked spiking and subthreshold responses of primary visual cortical (V1) neurons in awake mice presented with drifting grating visual stimuli of different orientations. For an orientation-selective neuron that was preferentially activated by gratings along specific orientations, we observed minimal voltage signal in response to non-preferred orientations but strong subthreshold as well as spiking activity in response to preferred orientations (**Fig. 3h**). From the spiking activity of all trials (ten 1-s trials for each of the eight gratings, **Fig. 3i**), we calculated the tuning curve of this neuron (black curve, **Fig. 3j**). Compared with the tuning curve calculated from subthreshold responses (red curve, **Fig. 3j**), the orientation tuning of spiking activity was sharper. Consistent with previous whole-cell recordings, these results suggest that spike thresholding sharpens selectivity for action potential output over subthreshold membrane potential [19, 20]. This same trend was observed in other orientation-as well as direction-selective neurons (**Supplementary Fig. 12**).

We also investigated how long it took for visual information to reach V1. In an example FOV, we found two visually responsive neurons at 345 μm below dura (**Fig. 3k**). The subthreshold responses reached their peaks 60 ms and 57 ms after the stimulus onset (i.e., the switch from dark screen to grating stimuli) (**Fig. 3l**), respectively. On the population level, the peak subthreshold response had a latency of 57.3 ± 2.3 ms (mean ± s.d., n = 20 cells from 3 mice, **Fig. 3m**), whereas the spike rate reached its peak 60 ms after stimulus onset (**Fig. 3n**). These latencies are consistent with the results measured with electrophysiological methods [21, 22].

In summary, using an all-optical passive laser scanner based on FACED, we achieved kilohertz-rate full-frame 2PFM imaging of neural activity with subcellular resolution in the mouse brain *in vivo*. FACED enabled electrical activity recordings of V1 neurons in awake mice, enabling characterization of orientation tuning properties and onset times of both subthreshold and spiking activity evoked by grating stimuli. The average laser power used in these experiments (**Supplementary Table 1**), 10–85 mW post objective, remained within the safe range and substantially below the threshold for heating-induced damages [23]. We did not observe signs of photodamages (e.g., blabbing of dendrites) in any of the samples tested, suggesting higher-order nonlinear photodamage processes to be also minimal. This is expected, because the power of individual beamlet post objective was ∼ 0.1–1 mW and overlaying tissue further reduced the actual focal energy as a result of scattering loss. Furthermore, subsequent excitation pulses arrived at the same sample positions after 1-ms delays, providing ample time for the fluorophores to return from their photodamage-prone dark states back to their ground electronic state, a process that was previously shown to reduce photobleaching and increase fluorophore brightness [24].

The frame rate of FACED 2PFM can be further increased by simply scanning the Y galvanometer at higher speeds. Using the 4-MHz 1,035-nm output of our laser system and scanning the Y galvanometer at 1,500 Hz, we imaged calcium activity *in vivo* at 3,000 fps with 80 × 1200 pixels per frame (**Supplementary Fig. 13, Supplementary Video 11**). New developments in laser systems (e.g., 4-MHz output at 920 nm, now commercially available) or voltage sensors (e.g., yellow or red variants that can be excited at 1,035 nm) should allow us to image voltage *in vivo* at 3,000 Hz as well, if desired. With the current pixel dwell time of 2 ns, FACED 2PFM reaches the limit on pixel rate imposed by the excited-state lifetime of high-quantum-yield fluorophores. Further reduction of the pixel dwell time would cause substantial cross-talk between neighboring pixels, resulting in a decrease in spatial resolution along the FACED axis. However, for activity imaging of spatially extended ROIs where signals from multiple pixels are averaged, minor cross-talk should not have a significant impact on activity measurement.

As we demonstrated here, by adding a passive FACED module to an existing 2PFM system, we transformed the microscope into an ultrafast imaging system with kHz frame rate. The FACED approach is thus readily compatible with conventional galvanometer-based 2PFMs, which favors its wide dissemination in optical brain imaging research. Following the conventional raster scanning strategy (albeit at much higher speeds), FACED 2PFM requires minimal computational processing, works in both sparsely and densely labeled samples, and is immune to crosstalk effects or source localization ambiguities observed in computation-based high-speed imaging techniques [12, 25]. Sampling an entire image plane allows post hoc motion correction, therefore making FACED 2PFM more resistant to sample motion than random-access 2PFMs. With existing sensors, FACED 2PFM offers sufficient speed and sensitivity to detect calcium and glutamate transients from neuronal processes, as well as both spiking and subthreshold voltage events from cell bodies. Future improvement in brightness and sensitivity of voltage indicators should further enhance detection of voltage signals in subcellular compartments, which should allow FACED imaging to fulfill its full potential in interrogating electrical activity in the brain at synaptic resolution.

## METHODS

### Animals

All animal experiments were conducted according to the National Institutes of Health guidelines for animal research. Procedures and protocols on mice were approved by the Institutional Animal Care and Use Committee at Janelia Research Campus, Howard Hughes Medical Institute.

### FACED two-photon fluorescence microscope (2PFM)

The simplified schematic of the FACED 2PFM is shown in **Supplementary Fig. 1a**. The two-photon excitation laser source at 920 nm (1 MHz repetition rate, 2 W maximal average output power, < 100 fs pulse width) was generated by an optical parametric amplifier (Opera-F, Coherent Inc.) that was pumped by a fiber laser at 1035 nm (Monaco 1035-40-40, 1 MHz or 4 MHz repetition rate, < 315 fs pulse width, 35 W, Coherent Inc.). After dispersion compensation [26], the laser beam was expanded with a 2× beam expander (BE02M-B, Thorlabs) to 8 mm in diameter. The beam then passed through a 4 to 5-mm wide slit and was one-dimensionally focused (Δθ = 1°) by a cylindrical lens (LJ1267RM-B, Thorlabs, effective input NA 0.01) into a nearly parallel mirror pair (*α* = 0.0125°, reflectivity > 99.9% and GDD<10 fs^2^ per reflection at 920 nm, fused silica substrate, 250 mm long and 20 mm wide, Layertec GmbH) with a separation of 300 mm. Because of the misalignment angle *α*, all the light rays eventually reflected back, following a set of zig-zig paths determined by their angles of incidence. After the FACED module, the rays (e.g. red rays in **Fig. 1a**) subjected to the same number of reflections by the mirror pair formed a single beamlet, and can be considered as emanating from a virtual light source located far away from the mirror pair. In this work, the retroreflected light rays formed 80 beamlets, and had their propagation distances within the FACED module (or their distance from their respective virtual sources) monotonically increase from ∼10 m to ∼60 m. The power throughput of the FACED module was ∼40%.

A polarization beam splitter (CCM1-PBS253, Thorlabs) in combination with a half-wave plate (AHWP05M-980, Thorlabs) and quarter-wave plate (AQWP05M-980, Thorlabs) were used to direct the spatially and temporally separated pulse trains into a 2PFM. A pair of achromatic doublets (AC508-500-B and AC508-250-B, Thorlabs) was used to conjugate the focal plan of the cylindrical lens and the midpoint of a pair of closely and orthogonally arranged X and Y galvo mirrors (6215H, Cambridge Technology).A scan lens / tube lens pair (SL50-2P2 and TTL200MP, Thorlabs) were used to conjugate the galvos to the back focal plane of a 25×/1.05 NA water-dipping objective lens (XLPLN25XWMP2, Olympus) that was mounted on a piezo stage (P-725K094, Physik Instrumente). The two-photon excited fluorescence signal was collected by the same microscope objective, reflected by a dichroic mirror (FF665-Di02-25×36, Semrock), focused by two lenses (AC508-080-A and LA1951-A Thorlabs), and after passing through an emission filter (FF01-680/SP, Semrock), detected by a photomultiplier tube without a current protection circuit (PMT, H7422P-40 MOD, Hamamatsu). The PMT signal was sampled at 625 MS/s with a high-speed digitizer (2G onboard memory, PXIe-5160, National Instruments) embedded in a chassis (PXIe-1071, National Instruments). From the chassis, the data was transferred to and saved by a desktop computer through a PCIe 16× interface.

By tuning the misalignment angle between the two mirrors (*α*= 0.0125°, see above), we generated a sequence of 80 focal spots extending 50 μm along the X axis. With the separation between the two mirrors set at 300 mm, the time delay between adjacent pulses was 2 ns. With 1 MHz repetition rate of the 920 nm output, the FACED module gave rise to a line scan rate of 1 MHz. Using the Y galvo to scan the foci along the direction orthogonal to the X/FACED axis at 500 Hz, and collecting the data bidirectionally, we achieved a frame rate of 1,000 fps with an effective image size of 80 × 900 pixels. 80 was given by the number of foci in the FACED axis and 900 was the product of the effective frame time (1 ms frame time minus a 100 μs dead time during mirror turns) and line scan rate. To increase the number of pixels in the FACED/X axis, X galvo was stepped to tile FACED images and increase the field of view. With 4 MHz 1035 nm excitation, we scanned the Y galvo at 1,500 Hz and achieved a frame rate of 3,000 Hz with an effective image size of 80 × 1200 pixels. For morphological imaging, we scanned the Y galvo at 50 Hz, resulting in a FACED imaging frame rate of 100 fps. (**Supplementary Table 1** listed all the major imaging parameters used in this work).

The raw data from the digitizer was saved as 1D waveforms. The Gen-1 PCIe 16× slot on the data acquisition computer had a maximal streaming rate of 250 Mb/s, which caused the data to overflow the on-chip memory of the digitizer after 6 s of data acquisition and thus limited the data collection to up to 6-s bouts. Upgrading the computer with Gen-3 PCIe 16× interface should allow us to stream data continuously. At 1 kHz frame rate, each image frame had 625 × 900 sampling points, with 100 × 900 data points sampling actual fluorescence excitation by the FACED foci and used to reconstruct a single image. If desired, multi-line scans were averaged to generate a single X line in the final image: for a Y-axis range of 150 μm, 3 line scans were averaged to form a single row; for a Y-axis range of 50 μm, 9 line scans are averaged. The final images were motion-registered with an iterative cross-correlation-based registration algorithm [27]. For morphological imaging, each FACED image frame had 625 × 9000 sampling points (10× exposure time), and a 10× increase in line averaging to form the final image.

### Analysis of activity data

We manually selected regions of interests (ROIs) from the averaged images of the registered image sequence. The mean fluorescent intensity within the ROIs was used to calculate the ΔF/F time traces, with F being the baseline fluorescence and ΔF being the fluorescence change due to neural activity. In the calcium and glutamate indicator datasets, to calculate the ΔF/F, we calculated the baseline fluorescence F by fitting the data points away from the transients (e.g. the first and last 1000 data points in **Fig. 2a**) with a single exponential function.

In the voltage indicator datasets, the ROI was selected to cover the cell membrane. Adjoining membrane segments between neighboring neurons were excluded from the ROI. A rolling percentile filter (50%, 500 ms window) was applied to the mean-intensity trace of the ROI to get the fluorescence baseline F (**Supplementary Fig. 9**) and rare periods with uncorrectable motion artifacts were discarded. In addition to ΔF/F traces, we calculated SNR traces as the ratio between ΔF (functional change) and √F (Poisson noise) [28]. For spike detection, the SNR traces were further subjected to a 250-Hz 12^th^ order low-pass Butterworth filter. The threshold for spikes was initially set at SNR = 7.5 and was manually adjusted if needed. Spikes were identified as local maxima that were at least 3 ms apart. To calculate the subthreshold activity, raw ΔF/F traces were low-pass filtered with a 50-Hz 12^th^ order low-pass Butterworth filter and the mean ΔF/F value during the entire duration of sensory stimulation (1-s of drifting grating) was used as the strength of subthreshold activity. To quantify the temporal dynamics of the optical voltage response, we aligned the optical spikes from the same neuron at peak response, and measured the rise (10% to 90%), decay (90% to 10%) and FWHM time from their averaged traces.

### Preparation and electric stimulation of primary neuronal culture

Primary neuronal cultures from neonatal rat pups were prepared as described previously [29]. AAV2/1.syn.GCaMP6s (1.8 × 10^13^ GC/ml), AAV.DJ.syn.iGluSnFR.A184V (3 × 10^12^ GC/ml), and AAV.DJ.syn.iGluSnFR.S72A (8.0 × 10^12^ GC/ml) were used to label cultured neurons by adding 1 μl of viral solution to each well in 24 well plates with 300 μl medium inside, respectively. After incubation overnight, 1 ml culture medium was added to each well. Neurons were imaged between 10 – 21 days post-transfection at room temperature in imaging buffer (145 mM NaCl, 2.5 mM KCl, 10 mM glucose, 10 mM HEPES, 2 mM CaCl_2_, 1 mM MgCl_2_, pH7.4).

For electric stimulation, cultured neurons in imaging buffer were positioned between two parallel electrodes separated at ∼10 mm. A stimulus isolator (NPIISO-01D100, ALA Scientific Instruments Inc.) and functional generator (AFG1022, Tektronix Inc.) was used to generate the electric field. For each stimulation, a train of 10 pulses (pulse duration: 1 ms; period: 12 ms; voltage: 50 V) was used to drive the neurons.

### Preparation and electric stimulation of acute brain slices

25-week-old male transgenic mice expressing GCaMP6f (scnn1a-TG3-cre x Ai93 x ACTB-tTA) [30, 31] were decapitated under deep isoflurane anesthesia, and the brain was transferred to an ice-cold dissection solution containing (in mM): 204.5 sucrose, 2.5 KCl, 1.25 NaH_2_PO_4_, 28 NaHCO_3_, 7 dextrose, 3 Na-pyruvate, 1 Na-ascorbate, 0.5 CaCl_2_, 7 MgCl_2_ (pH 7.4, oxygenated with 95% CO_2_ and 5% O_2_). 350-μm-thick coronal slices of the primary visual cortex (V1) were sectioned using a vibrating tissue slicer (Leica VT 1200S, Leica Microsystems, Wetzlar, Germany). The slices were then transferred to a suspended mesh within an incubation chamber filled with artificial cerebrospinal fluid (ACSF) containing (in mM): 125 NaCl, 2.5 KCl, 1.25 NaH_2_PO_4_, 25 NaHCO_3_, 25 dextrose, 1.3 CaCl_2_, 1 MgCl_2_ (pH 7.4, oxygenated with 95% CO_2_ and 5% O_2_). After 30 – 60 minutes of recovery at 35°C, the chamber was maintained at room temperature.

During imaging, slices were submerged in a recording chamber constantly perfused with oxygenated ACSF. A micropipette filled with ACSF were used for monopolar stimulation via a stimulus isolator (NPIISO-01D100, ALA Scientific Instruments Inc.) and a function generator (AFG1022, Tektronix Inc.). To provide extracellular stimulation, the stimulating electrode was placed in the proximity of the recorded cell, and a train of 10 pulses (pulse duration: 1 ms; period: 12 ms; current: 300 μA) was applied.

### Mouse preparation for *in vivo* imaging

Mice (females or males, >2-months-old) were housed in cages (in groups of 1 – 5 before surgeries and in pairs or single housed after) under reverse light cycle. Wild-type (Jackson Laboratories, Black 6, stock #:000664) as well as Gad2-IRES-cre (Jackson Laboratories, Gad2tm2 (cre) Zjh/J, stock #: 010802) mice were used.

Virus injection and cranial window implantation procedures have been described previously [22]. Briefly, mice were anaesthetized with isoflurane (1 – 2% by volume in O_2_) and given the analgesic buprenorphine (SC, 0.3 mg per kg of body weight). Animals were head fixed in a stereotaxic apparatus (Model 1900, David Kopf Instruments). A 3.5-mm diameter craniotomy was made over the left V1 with dura left intact. A glass pipette (Drummond Scientific Company) beveled at 45° with a 15 – 20 μm opening was back-filled with mineral oil. A fitted plunger controlled by a hydraulic manipulator (Narishige, MO10) was inserted into the pipette and used to load and slowly inject 30 nl viral solution into the brain (∼200 – 400 μm below pia). 3 – 6 injection sites were chosen in the left V1 with 0.3 to 0.5 mm space between injection sites. The following viral vectors were used to label neurons with different sensors. Labeling with calcium sensor: AAV2/1.syn.GCaMP6s (1.8 × 10^13^ GC/ml); Dense labeling with glutamate sensors: AAV.DJ.syn.iGluSnFR.A184V (3 × 10^12^ GC/ml); AAV.DJ.syn.iGluSnFR.S72A (8.0 × 10^12^ GC/ml); Sparse labeling with glutamate sensors: AAV.DJ.syn.FLEX.iGluSnFR.A184V (2.8 × 10^13^ GC/ml) 1:1 mixed with AAV2/1.syn.Cre (500 times diluted from 1.5 × 10^13^ GC/ml); AAV.DJ.syn.FLEX.iGluSnFR.S72A (5.2 × 10^12^ GC/ml) 1:1 mixed with AAV2/1.syn.Cre (500 times diluted from 1.5 × 10^13^ GC/ml); labeling with the voltage sensor: AAV2/9.syn.ASAP3-Kv (1.55 × 10^12^ GC/ml). At the completion of viral injections, a glass window made of a single coverslip (Fisher Scientific No. 1.5) was embedded in the craniotomy and sealed in place with dental acrylic. A titanium head-post was then attached to the skull with cyanoacrylate glue and dental acrylic. *In vivo* imaging was carried out after at least two weeks of recovery with single or paired housing and habituation for head fixation. All imaging experiments were carried out on head-fixed awake mice.

### Visual stimulation in head-fixed awake mice

Visual stimuli were presented by a liquid crystal display (22-inch diagonal and 1920 × 1080 pixels). The screen was positioned at 15 cm from the eye of the mice and orientated at ∼40° to the long axis of the mice. Drifting sinusoidal gratings were presented for 1.5 s at 8 orientations (0° to 315° at 45° steps) in pseudorandom sequences. Between the grating stimulus, 6s dark screen were presented. Gratings had 100% contrast and 0.06 cycle per degree and drifted at 2 Hz. During each 1.5 s stimulation period, a sequence of 1,000 images were recorded from 0 s to 1 s; during each 6 s dark adaptation period, a sequence of 1,000 images were recorded from 2.5 s to 3.5 s. A total of 5 or 10 trials were repeated for each stimulus.

### Data processing

Unless stated otherwise, all images and data presented here were unprocessed raw images/data, without smoothing, denoising, or deconvolution. **Supplementary Videos 1-10** were collected at 1,000 fps but were binned every 20 or 50 frames (no binning in **Supplementary Video 10**) and saved at 20 binned fps for video output by Fiji [32], which was not capable of saving videos at 1,000 fps. **Supplementary Video 11** were collected at 3,000 fps but were binned every 60 frames and saved at 20 binned fps for video output. “Green hot” lookup table in ImageJ was used for all images.

## ACKNOWLEDGEMENTS

The authors thank Cristina Rodriguez for help with the laser system; Rongwen Lu for help with visual stimulation experiments; Guan Cao for providing cultured neuron samples; Jonathan Marvin, Loren Looger for providing glutamate sensors; and the Janelia JET team for designing and assembling the dispersion compensation unit. This work was supported by Howard Hughes Medical Institute (J.W., Y.L., C.-L.H., N.J.), and American Epilepsy Society predoctoral fellowship (M.C.); the China Scholarship Council Joint PhD Training Program (D.S.); Stanford Neuroscience PhD Program training grant 5T32MH020016 and the Post-9/11 GI Bill (S.W.E.); Research Grants Council of the Hong Kong Special Administrative Region of China (17209017, 17259316, 17207715) (J.W., K.K.T.); and NIH BRAIN Initiative grants 1U01NS103464 (M.Z.L.), 1RF1MH114105 (M.Z.L.), and 1UF1NS107696 (J.W., N.J., K.K.T.).

## CONTRIBUTIONS

NJ conceived of the project; ML, KKT, and NJ supervised research; JW, KKT, and NJ designed FACED module; YL, SC, CLH prepared samples; MC created ASAP3; MC, SE, and DS characterized ASAP3 and ASAP3-expressing viruses; JW collected and analyzed the data; JW and NJ wrote the manuscript with inputs from all authors.

## SUPPLEMENTARY VIDEO INFORAMTION

**Supplementary video 1: Imaging calcium transients at 1,000 fps in GCaMP6s-expressing cultured neurons evoked by extracellular electric stimulation.** Same data as in Supp. Fig. 3. The raw image sequence was binned every 50 frames, saved at 20 binned fps, and compressed for video output.

**Supplementary video 2: Imaging calcium transients at 1,000 fps in GCaMP6f-expressing neurons in acute mouse brain slices evoked by extracellular electric stimulation.** Same data as in Supp. Fig. 4. The raw image sequence was binned every 50 frames, saved at 20 binned fps, and compressed for video output.

**Supplementary video 3: Imaging spontaneous calcium release events at 1,000 fps in neurites of GCaMP6s-expressing cultured neurons.** Same data as in Supp. Fig. 5. The raw image sequence was binned every 20 frames, saved at 20 binned fps, and compressed for video output.

**Supplementary videos 4-5: Imaging glutamate transients at 1,000 fps in cultured neurons expressing A184V variant of iGluSnFR evoked by extracellular electric stimulation.** Same A184V data as in Fig. 2a. The raw image sequence was binned every 50 frames, saved at 20 binned fps, and compressed for video output.

**Supplementary videos 6-7: Imaging glutamate transients at 1,000 fps in cultured neurons expressing S72A variant of iGluSnFR evoked by extracellular electric stimulation.** Same S72A data as in Fig. 2a. The raw image sequence was binned every 50 frames, saved at 20 binned fps, and compressed for video output.

**Supplementary video 8: Imaging glutamate transients of layer 2/3 neurons expressing A184V variant of iGluSnFR in V1 of awake mice at 1,000 fps.** The white arrow points to the glutamate releasing site. Same A184V data as in Fig. 2b. The raw image sequence was binned every 20 frames, saved at 20 binned fps, and compressed for video output.

**Supplementary video 9: Imaging glutamate transients of layer 2/3 neurons expressing S72A variant of iGluSnFR in V1 of awake mice at 1,000 fps.** The white arrow points to the glutamate releasing site. Same S72A data as in Fig. 2b. The raw image sequence was binned every 20 frames, saved at 20 binned fps, and compressed for video output.

**Supplementary video 10: Imaging rapid movement of a fluorescent particle at 1,000 fps in the layer 2/3 of awake mouse brain** *in vivo*. Same data as in Supp. Fig. 6. A sequence of 50 raw images recorded at 1,000 fps was saved at 20 fps and compressed for video output.

**Supplementary video 11: Imaging spontaneous calcium transients of layer 2/3 neurons expressing GCaMP6s in V1 of awake mice at 3,000 fps.** Same data as in Supp. Fig. 14. The raw image sequence was binned every 60 frames, saved at 20 binned fps, and compressed for video output.

---

**Supplementary Figure 1:**
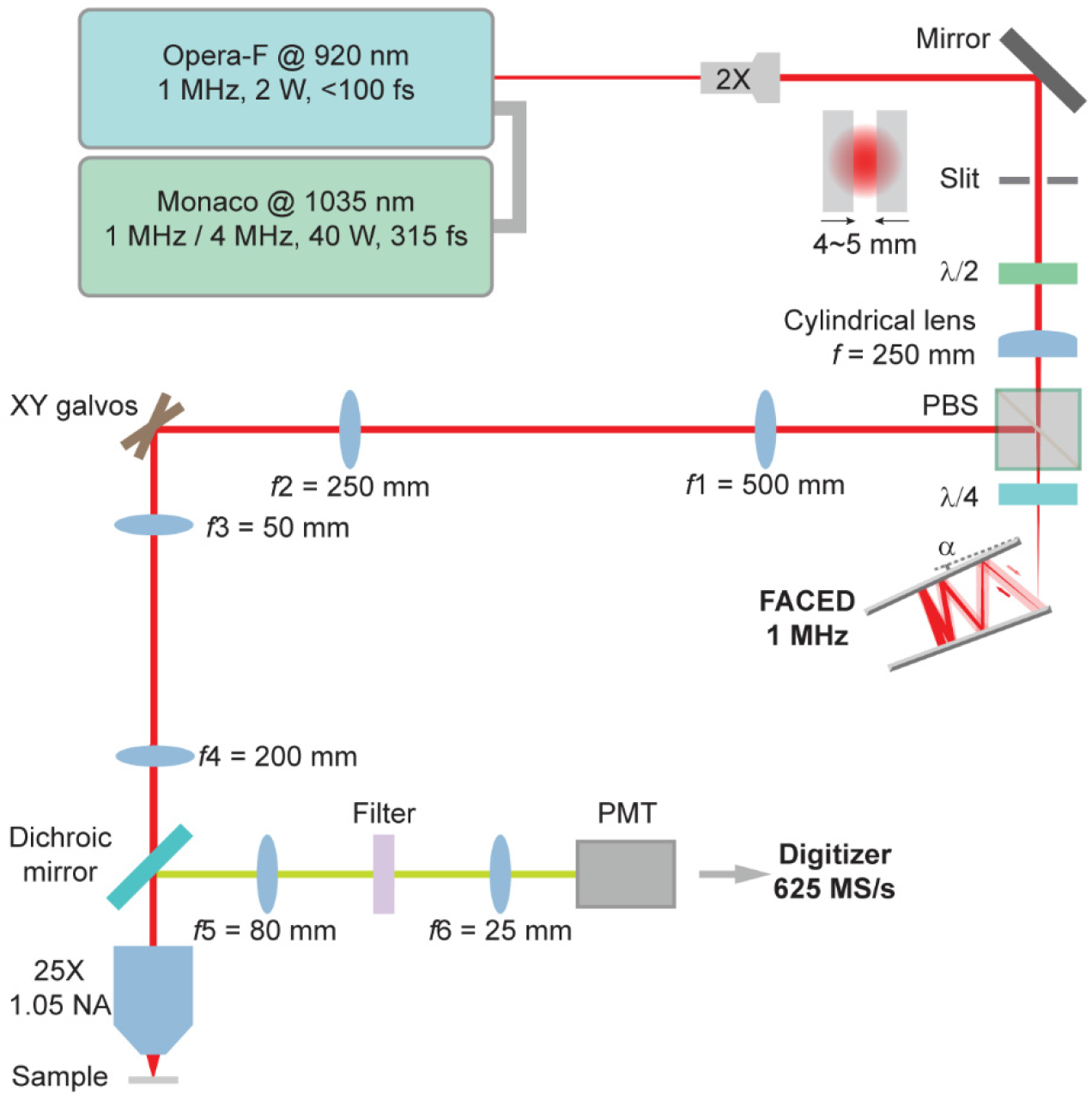
Optical layout of the FACED two-photon fluorescence microscope. 2X: 2-fold beam expander; λ/2: Half-wave plate; PBS: polarizing beam splitter; λ/4: quarter-wave plate; α: misalignment angle of the mirror pair; PMT: photomultiplier tube. The FACED module scans the foci along the X galvo direction. Note that the X galvo is deactivated for high-speed functional imaging.

**Supplementary Figure 2:**
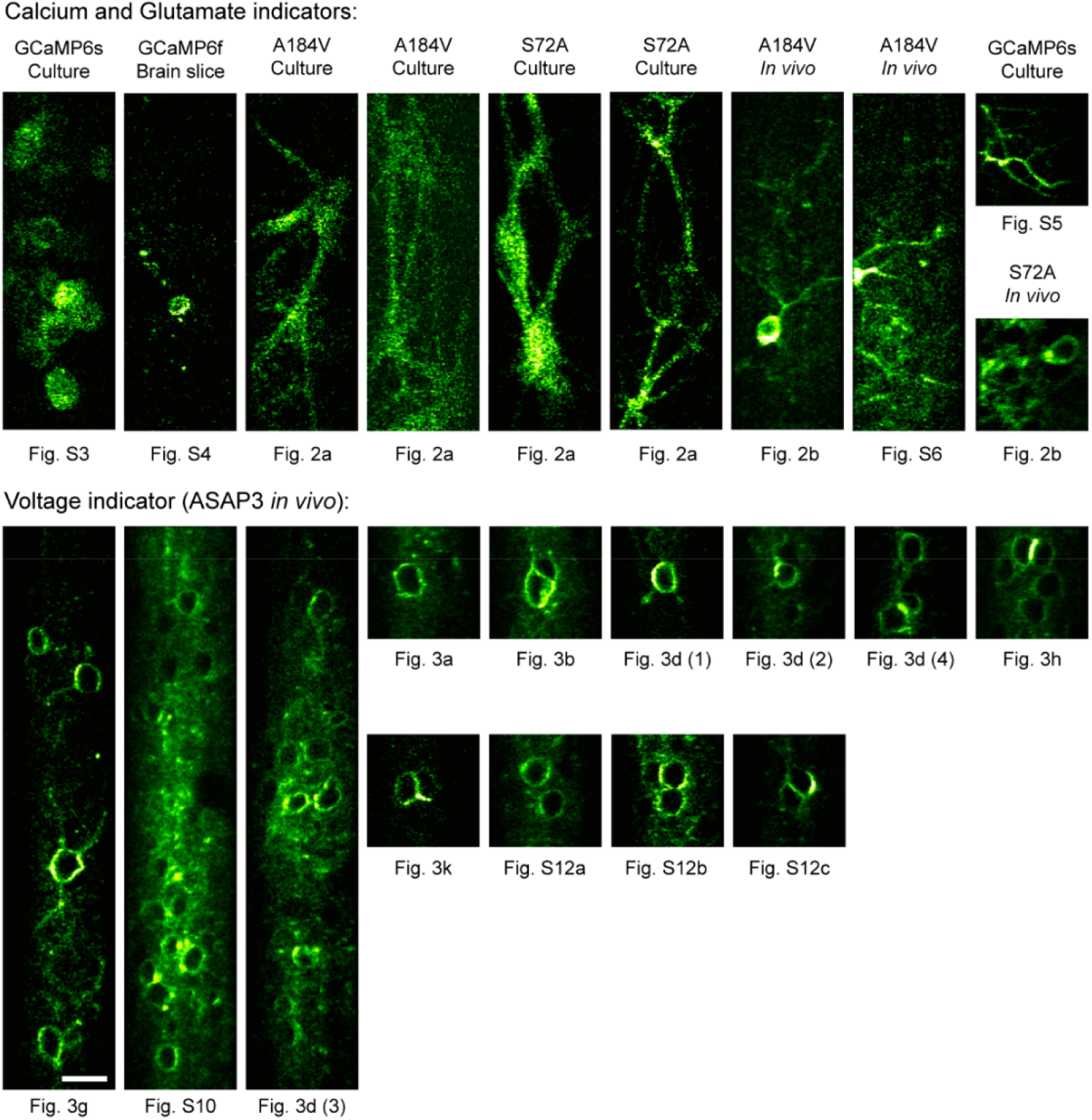
Representative raw images taken at 1,000 fps for different indicators. Calcium indicators: GCaMP6s/6f; Glutamate indicators: A184V and S72A variants of SF-iGluSnFR; voltage indicator: ASAP3-Kv. Scale bar: 20 μm.

**Supplementary Figure 3:**
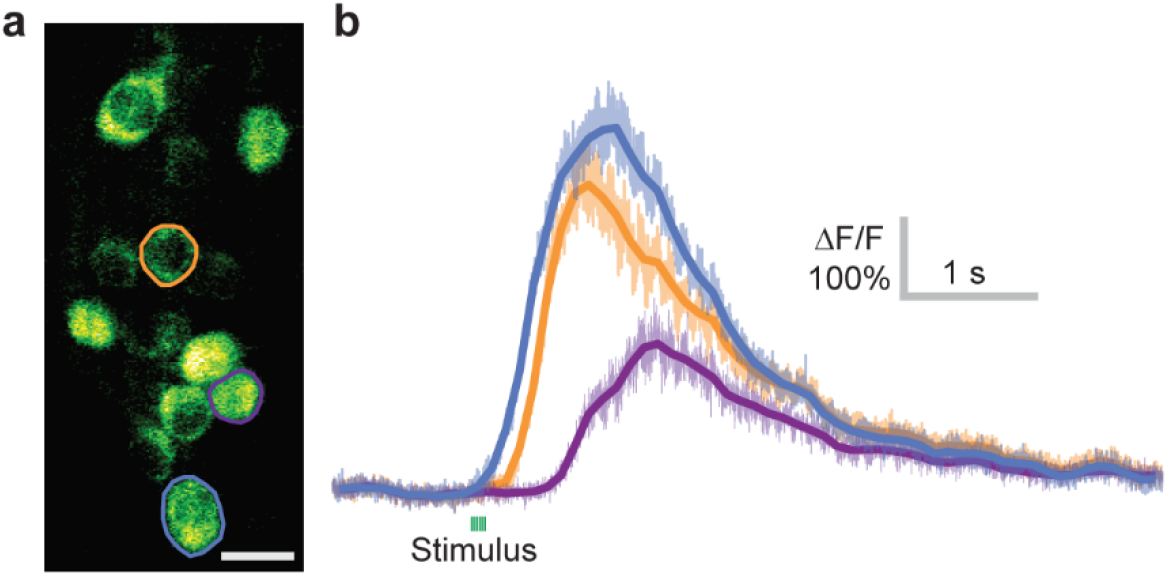
1 kHz imaging of GCaMP6s-expressing cultured neurons. (a) Morphological image. (b) Calcium transients of 3 cells in (a) evoked by field electrode stimulation and recorded at 1,000 fps. Darker lines were 100-point boxcar averages of the raw traces to guide the eye. Scale bar: 20 μm.

**Supplementary Figure 4:**
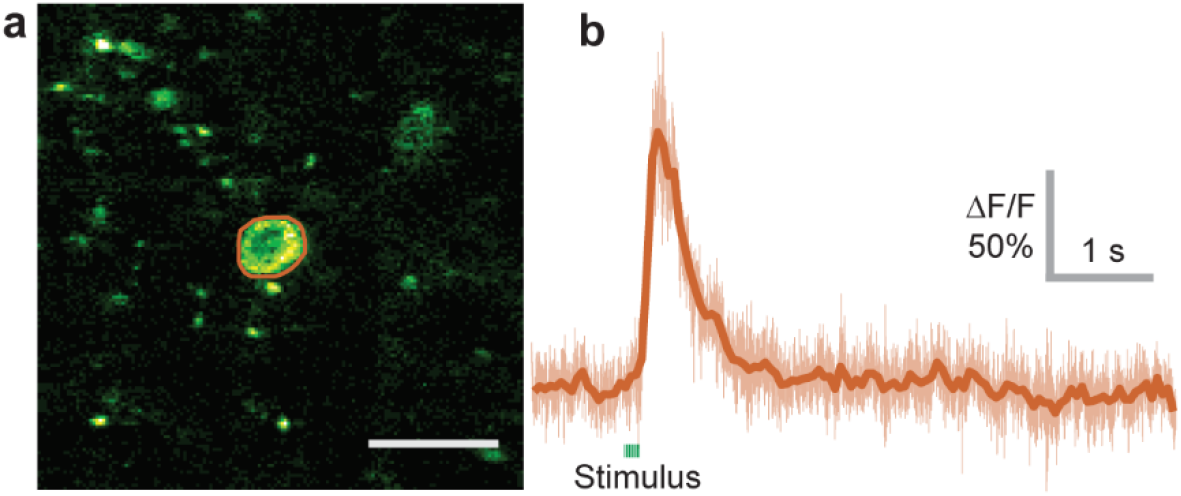
1 kHz imaging of a GCaMP6f-expressing neuron in an acute brain slice. (a) Morphological image. (b) Calcium transient of the cell in (a) evoked by field electrode stimulation and recorded at 1,000 fps. Darker line was 50-point boxcar average of the raw trace to guide the eye. Scale bar: 20 μm.

**Supplementary Figure 5:**
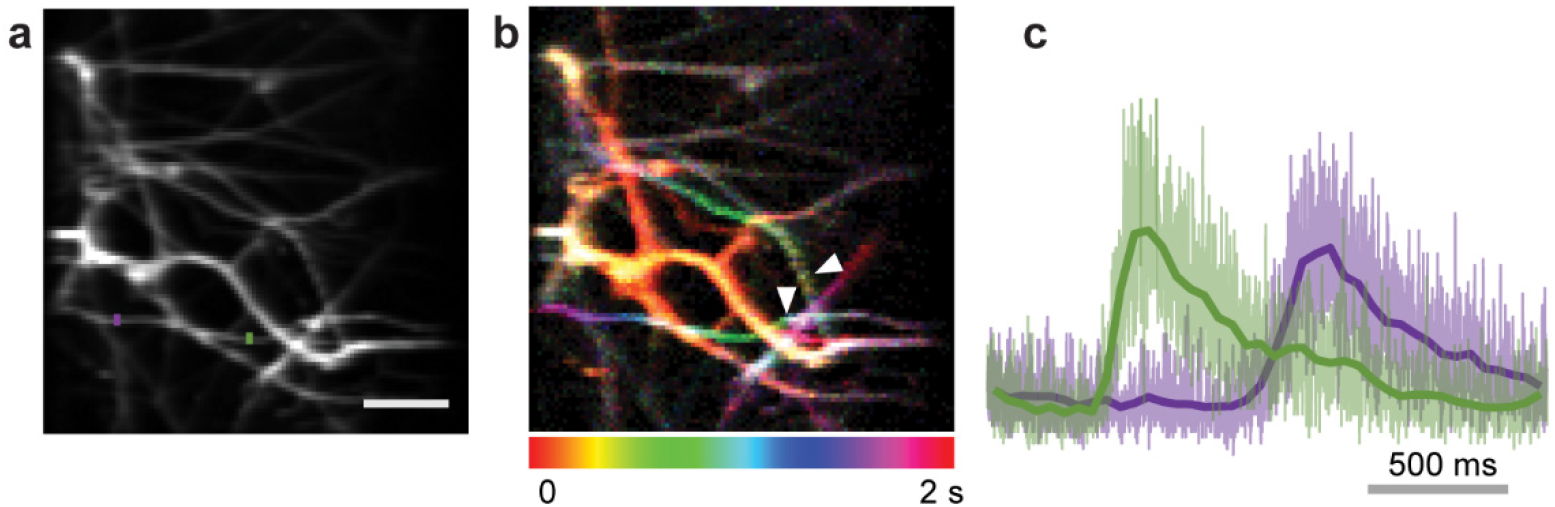
1 kHz imaging of spontaneous calcium increases in neurites of GCaMP6s-expressing cultured neurons. (a) Mean intensity projection of 2,000 frames. (b) Temporal color coding of the 2,000 frames highlights the sites where calcium increases were initially observed (white arrowheads). (c) Calcium transients at the color masked positions in (a), indicating a calcium propagating speed of ∼25 μm/s. Darker lines were 50-point boxcar averages of the raw traces to guide the eye. Scale bar: 10 μm.

**Supplementary Figure 6:**
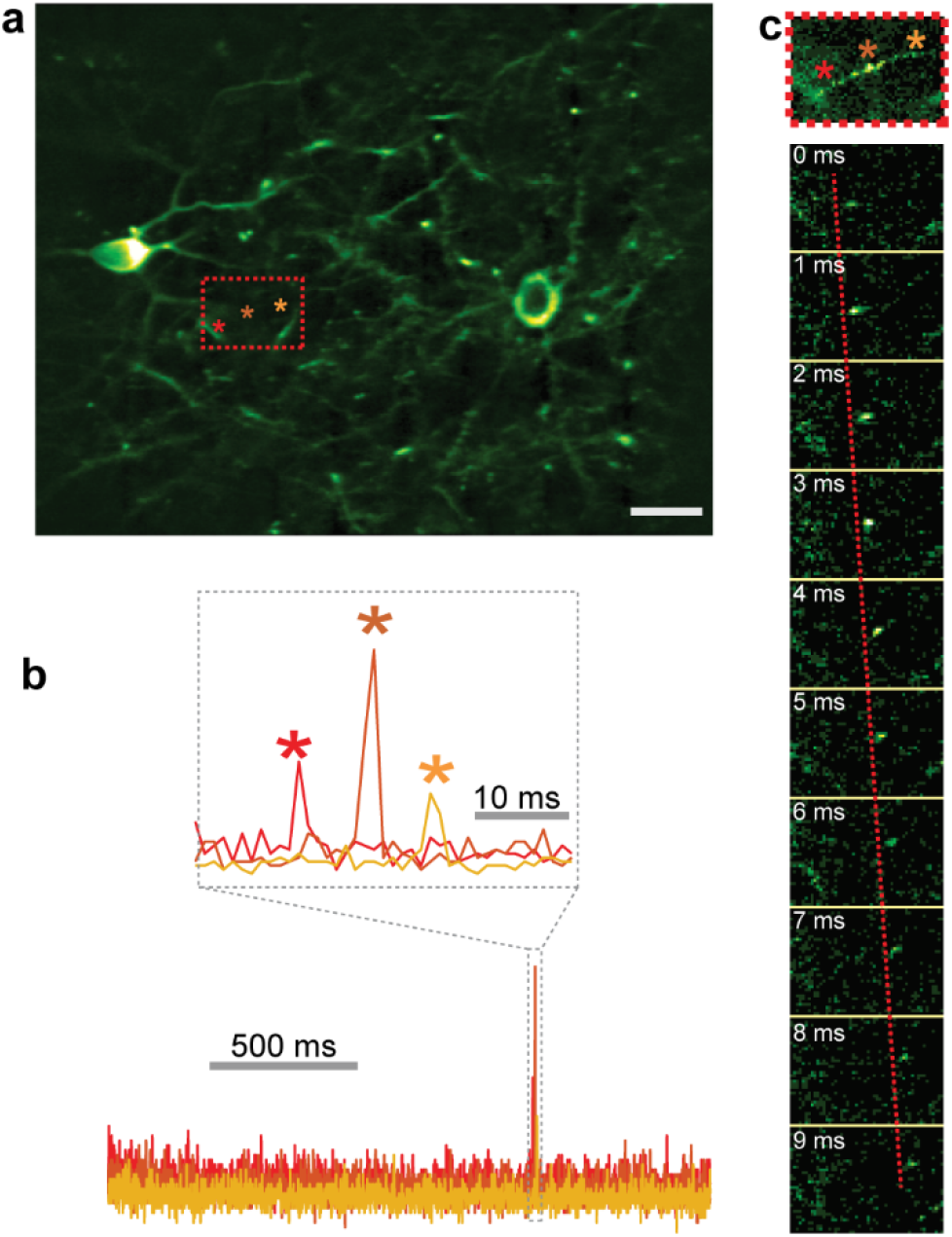
Tracking rapid movement of a fluorescent particle in the awake mouse brain *in vivo*. (a) Morphological image of layer-2/3 neurons expressing the A184V variant of the glutamate sensor iGluSnFR in V1 of an awake mouse. (b) sssTime traces at the three pixels indicated by the three asterisks in (a). (c) Top panel shows the maximum intensity projection of the 10 image frames collected at 1 kHz of the boxed region in (a); bottom panel shows the ten individual frames imaged at 1-ms intervals. Scale bars: 20 μm.

**Supplementary Figure 7:**
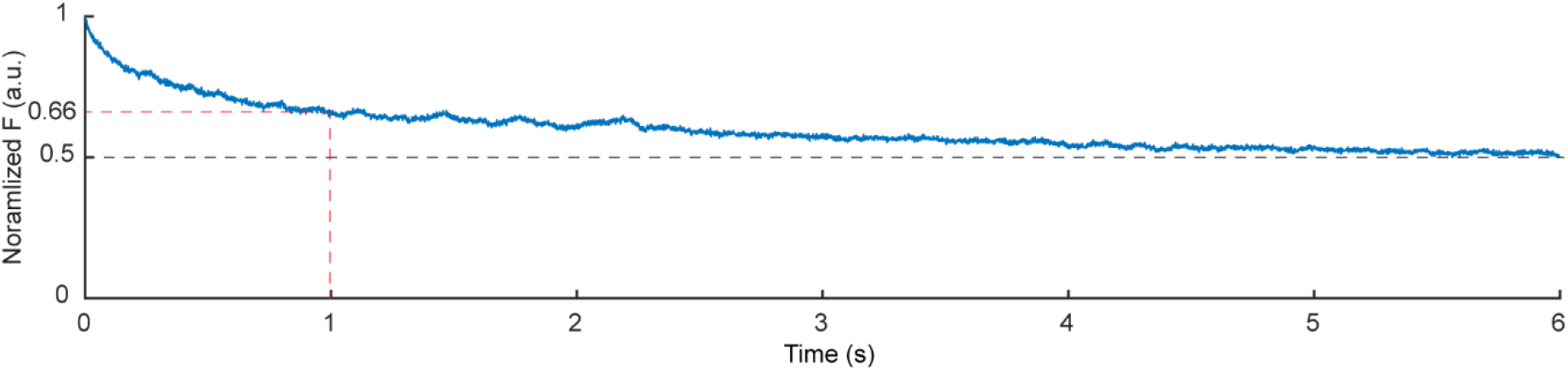
Representative ASAP3-Kv fluorescence photobleaching during 6-s continuous 1KHz FACED 2PFM imaging. The mean fluoresce intensity for the field of view (FOV) in **Fig. 3g** during a 6-sec recording. Imaging depth: 75 μm; FOV: 50×250 μm^2^; post-objective power: 35 mW.

**Supplementary Figure 8:**
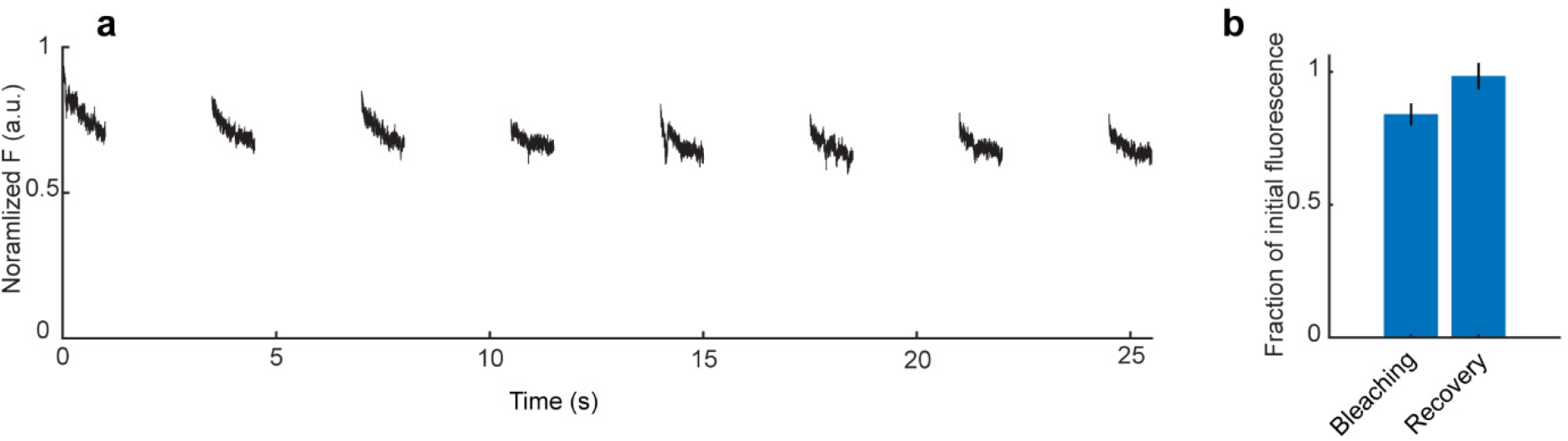
Representative bleaching and recovery dynamics of ASAP3-Kv fluorescence during intermittent 1-s duration 1KHz FACED 2PFM imaging. Data obtained from ROI 2 in Fig. 3b. (a) Fluorescence signal change for eight 1-s recordings that were collected with a 2.5-s gap in between. (b) Histogram of the bleaching and recovery level from 80 1-s 1,000-fps recordings. Bleaching is quantified by the ratio between the mean fluorescence signal of the last 50 frames and first 50 frames within a sequence of 1,000 frames (0.84 ± 0.04). Recovery is quantified by the ratio of mean fluorescence signal between the n+1^th^ 1,000 frames and n^th^ 1,000 frames (0.98 ± 0.05). Imaging depth: 125 μm; FOV: 50×50 μm^2^; post-objective power: 35 mW.

**Supplementary Figure 9:**
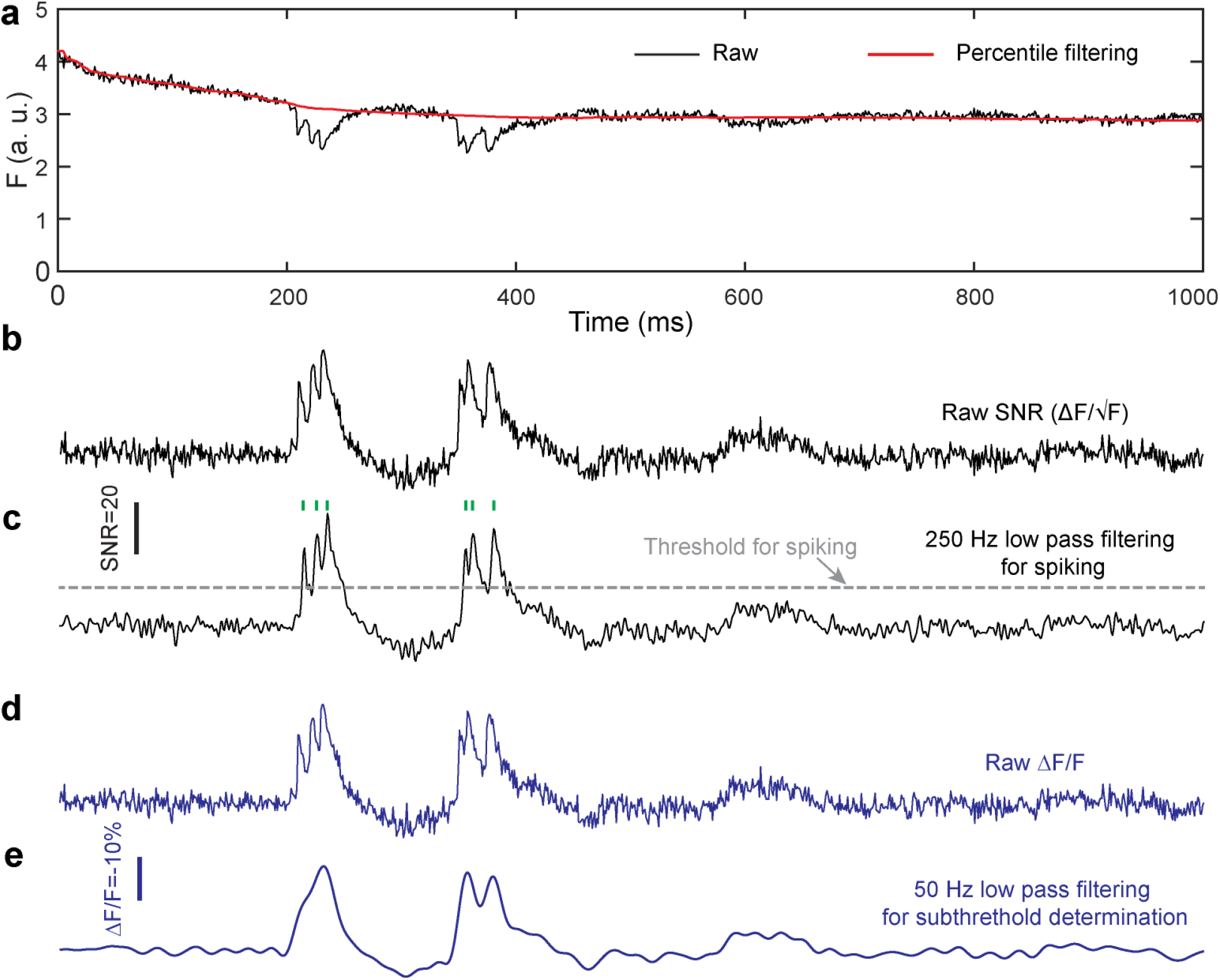
Data processing for *in vivo* voltage traces. (a) Baseline fluorescence F (red) was obtained by low-pass filtering (500-ms rolling percentile (50%) filter) of the raw signal trace (black). (b) Raw SNR trace (ΔF/√F, fluorescence change-to-noise ratio). (c) Smoothened SNR trace obtained by applying a 250-Hz low-pass Butterworth filter to (b). Optical spikes (green ticks) were identified as local SNR maxima above a spiking threshold (e.g., 7.5) at least 3 ms apart. (d) Raw ΔF/F. (e) Smoothened trace representing the subthreshold response obtained by applying a 50-Hz low-pass Butterworth filter to (d). All traces in **Fig. 3** are raw traces as in (b) and (d).

**Supplementary Figure 10:**
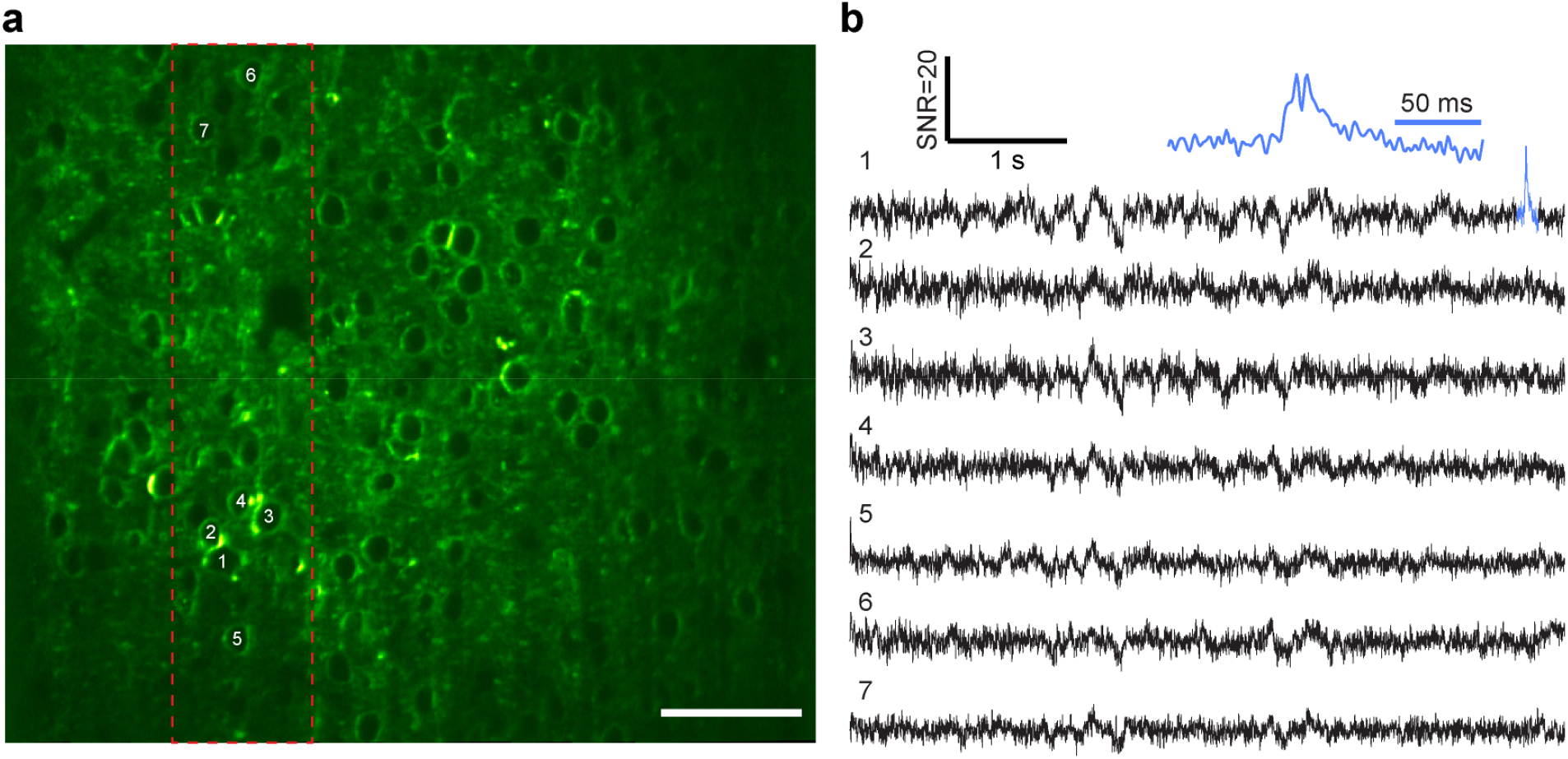
1 kHz imaging of voltage responses in V1 of awake mice at a 50 × 250 μm^2^ FOV. (a) Dashed red box indicates the FOV for 1kHz FACED imaging; within this FOV, 15 ASAP3-expressing cells were imaged simultaneously. (b) Voltage traces for the 7 cells labeled in (a). During a 6-s recording, a burst of two spikes were detected for ROI 1. Note the absence of these spikes in ROIs 2-5, which were in close proximity with ROI 1. Image depth: 155 μm; post-objective power: 75 mW. Scale bar: 50 μm.

**Supplementary Figure 11:**
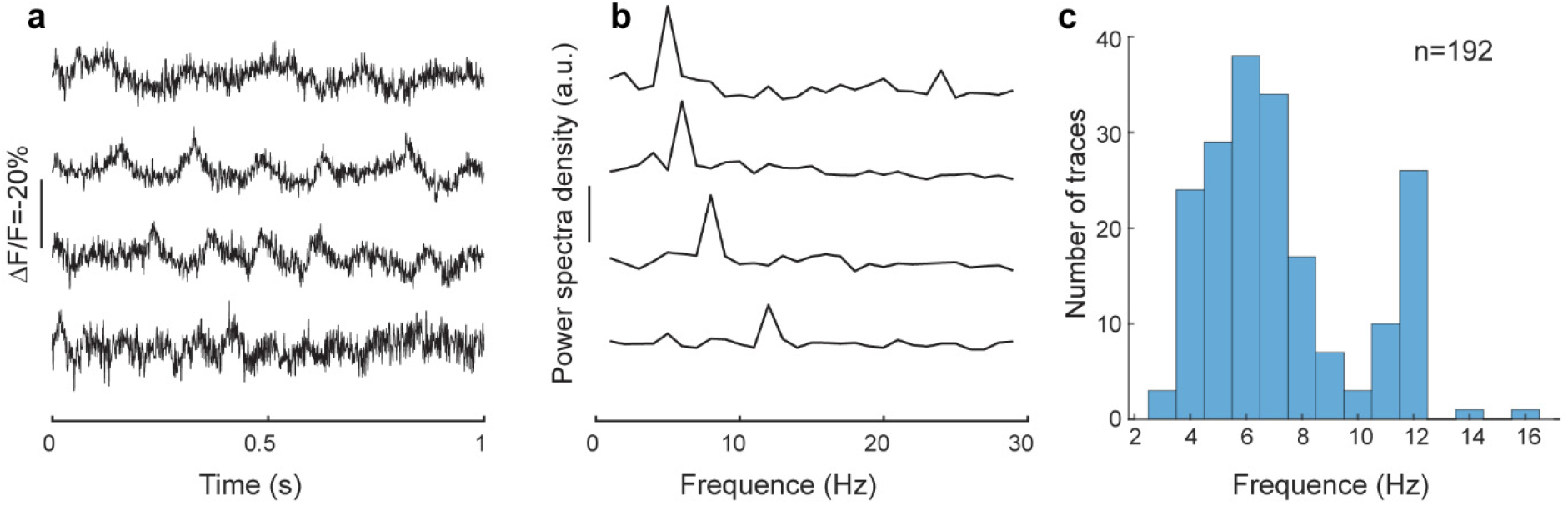
Imaging the subthreshold oscillations in V1 of awake mice. (a) Representative raw ΔF/F traces and (b) their corresponding power spectra from individual neurons. (c) Histogram of the peak oscillation frequencies (192 traces from 120 cells in three mice).

**Supplementary Figure 12:**
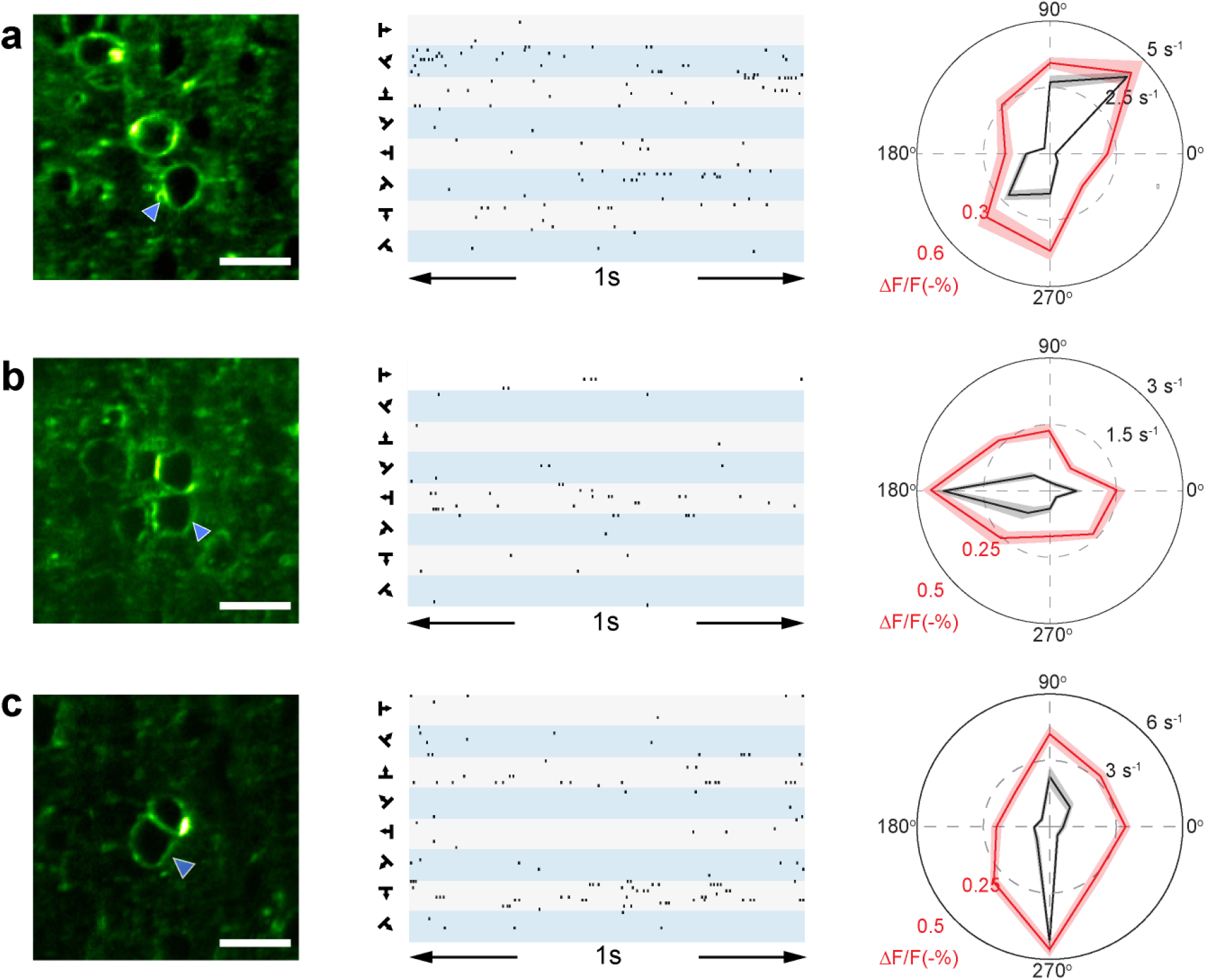
Orientation and direction tuning of neurons in V1 of awake mice. Left panels: FACED 2PFM images of the neurons. Middle panels: Raster plots showing optical spikes detected from each neuron (blue arrowheads) during the presentation of grating stimuli (10 trials for each of the 8 stimuli). Right panels: Polar plots showing mean spike rate (black) and average subthreshold ΔF/F (red) for (a) an orientation-selective and (b,c) two direction-selective neurons. Shaded areas: standard deviation. Image depth and post-objective power for a, b, c: 140, 170, 130 μm; 35, 30, 30 mW. Scale bar: 20 μm.

**Supplementary figure 13:**
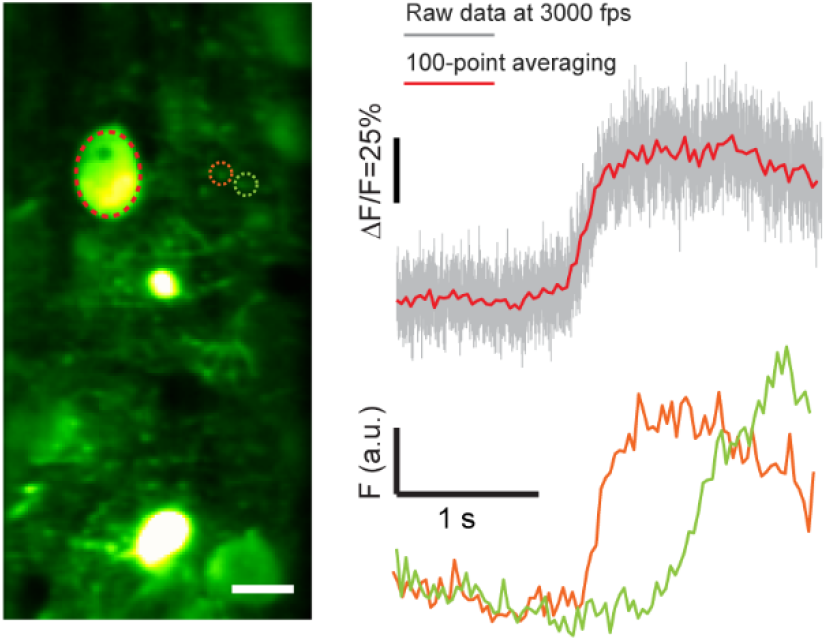
3 kHz neural activity imaging in V1 of awake mouse *in vivo*. (Left) Mean intensity projection of 9,000 frames recorded in 3 s; (Right) Top panel: a spontaneous calcium transient from the soma of a cell (red dashed ROI); bottom panel: fluorescence traces within two ROIs (orange and green ROIs) show calcium propagation along a dendrite. Gray lines: raw data traces; colored lines: 100-point boxcar averaging of the raw traces. Indicator: GCaMP6s; imaging depth: 125 μm; FOV: 50×100 μm^2^; post-objective power: 90 mW; excitation wavelength: 1035 nm.

**Supplementary table 1:**
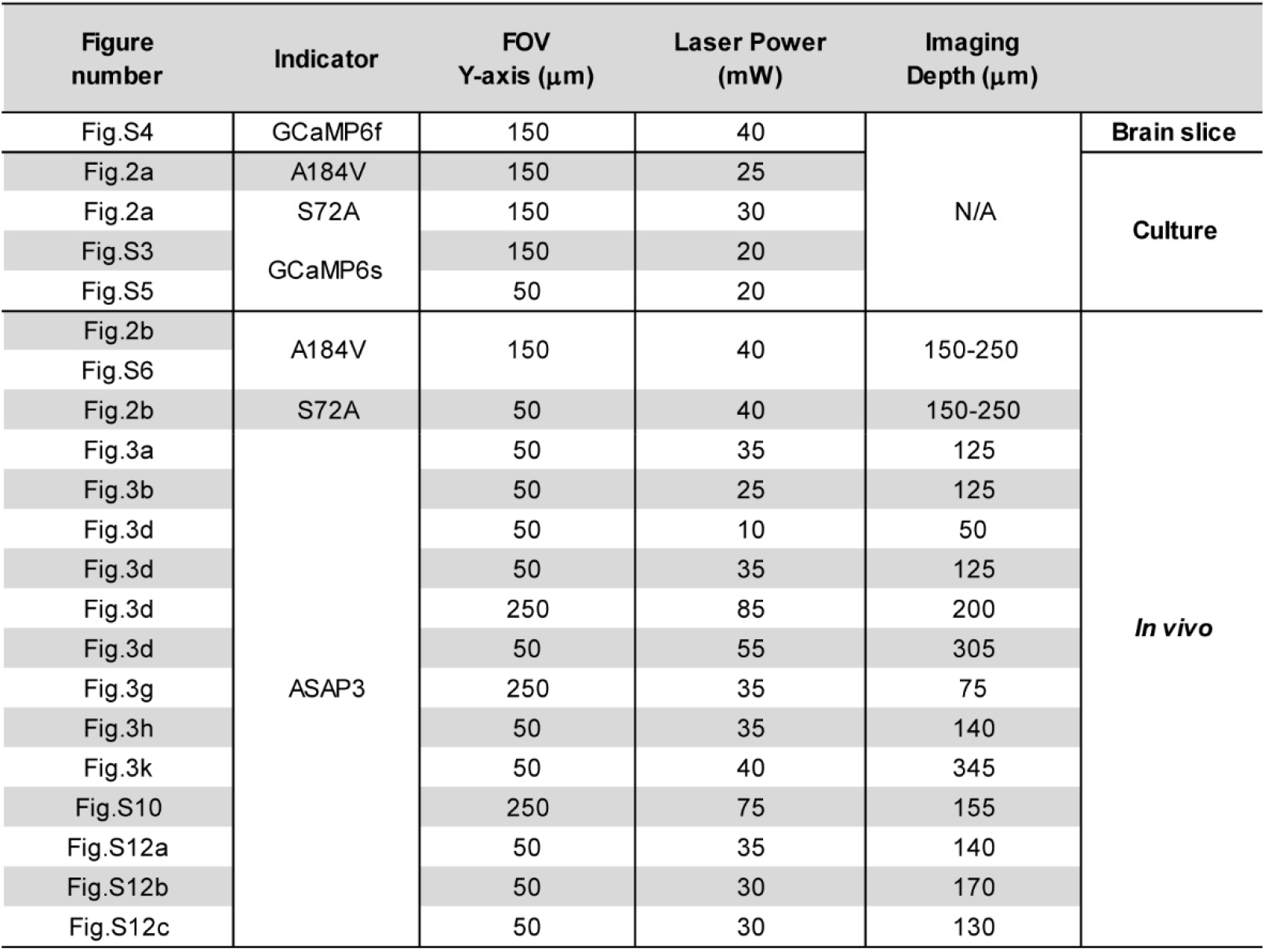
Major parameters used for FACED imaging at 1 kHz

